# Multi-scale periodicities in the functional brain networks of patients with epilepsy and their effect on seizure detection

**DOI:** 10.1101/221036

**Authors:** Georgios D. Mitsis, Maria Anastasiadou, Manolis Christodoulakis, Eleftherios S. Papathanasiou, Savvas S. Papacostas, Avgis Hadjipapas

## Abstract

The task of automated epileptic seizure detection and prediction by using non-invasive measurements such as scalp EEG signals or invasive, intracranial recordings, has been at the heart of epilepsy studies for at least three decades. By far, the most common approach for tackling this problem is to examine short-length recordings around the occurrence of a seizure - normally ranging between several seconds and up to a few minutes before and after the epileptic event - and identify any significant changes that occur before or during the event. An inherent assumption in these studies is the presence of a relatively constant EEG activity in the interictal period, which is presumably interrupted by the occurrence of a seizure. Here, we examine this assumption by using long-duration scalp EEG data (ranging between 21 and 94 hours) in patients with epilepsy, based on which we construct functional brain networks. Our results suggest that not only these networks vary over time, but they do so in a periodic fashion, exhibiting multiple peaks at periods ranging between around one and 24 hours. The effects of seizure onset on the functional brain network properties were found to be considerably smaller in magnitude compared to the changes due to the inherent periodic cycles of these networks. Importantly, the properties of the identified network periodic components (instantaneous phase, particularly that of short-term periodicities around 3 and 5 h) were found to be strongly correlated to seizure onset. These correlations were found to be largely absent between EEG signal periodicities and seizure onset, suggesting that higher specificity may be achieved by using network-based metrics. In turn, this suggests that to achieve more robust seizure detection and/or prediction, the evolution of the underlying longer term functional brain network periodic variations should be taken into account.

**Highlights:** - We have examined the long-term characteristics of EEG functional brain networks and their correlations to seizure onset

- We show periodicities over multiple time scales in network summative properties (degree, efficiency, clustering coefficient)

- We also show that, in addition to average network properties, similar periodicities exist in network topology using a novel measure based on the graph edit distance, suggesting that specific connectivity patterns recur over time

- These periodic patterns were preserved when we corrected for the effects of volume conduction and were found to be of much larger magnitude compared to seizure-induced modulations

- For the first time to our knowledge, we demonstrate that seizure onset occurs preferentially at specific phases of network periodic components that were consistently observed across subjects, particularly for shorter periodicities (around 3 and 5 hours)

- These correlations between the phase of network periodic components and seizure onset were nearly absent when examining univariate properties (EEG signal power), suggesting that network-based measures are more tightly coupled with seizure onset compared to EEG signal-based measures

- Our findings suggest that seizure detection and prediction algorithms may benefit significantly by taking into account longer-term variations in brain network properties

- As we show strong evidence that shorter network-based periodicities (3-5 hours) are tightly coupled with seizure onset, our results pave the way for further investigation into the pathophysiology of seizure generation mechanisms beyond the well-known effects of circadian rhythms

## 1 Introduction

The task of detecting or predicting epileptic seizures has received tremendous attention for more than 30 years. Automated detection and prediction algorithms based on electroencephalographic (EEG) measurements attempt to characterize the transition from the inter-ictal to the ictal state, by identifying EEG patterns that significantly deviate from the interictal state. For this reason, knowledge of the baseline inter-ictal properties is vital. However, an inherent assumption commonly made is that EEG activity during this inter-ictal state is relatively constant and interrupted by seizure occurrence. This assumption arises, at least in part, because only short-length recordings (several seconds to minutes) around seizure onset are typically examined.

However, assuming a constant baseline (inter-ictal state) is at odds with the long-established influence of longer term biological rhythms (e.g. the circadian rhythm) on physiological signals (Glass 2001). These signals include heartbeat dynamics and heart rate variability, blood pressure and EEG among others and typically exhibit a 1/f behavior in the frequency domain. Specifically, the influence of the circadian rhythm on EEG signal properties, which results approximately in a main 24-hour periodicity, has been demonstrated for almost half a century (Scheich 1969). Kaiser et al. (Kaiser and Sterman 1994; Kaiser 2008) studied healthy subjects for twelve hours during the daytime wakefulness and identified a weak ultradian modulation with a cycle of approximately 90-120 min in the 9-11 Hz frequency band, and a slower and stronger temporal modulation with 4 hours period in the 11-13 Hz band. A strong 3-4-hour periodicity in the EEG power during daytime, which was most pronounced in the fronto-central frequencies over 22.5 Hz, as well as in parietal alpha activity, was reported in (Chapotot et al. (2000)).

Some studies related to EEG-based epilepsy detection and prediction have also demonstrated the effect of the circadian rhythm. For instance, (Kreuz et al. (2004)) studied one patient suffering from epilepsy and observed that circadian patterns were clearly observable in certain intracranial channel combinations. (Schad et al. (2008)) observed, in a subset of their patient population, an effect of the circadian rhythm on features obtained from the spiking rate, i.e. the rate at which local slopes cross a specified threshold, of both scalp and intracranial EEG. More recent studies have suggested that seizures tend to occur at specific times during the day and that this information can be used to improve seizure prediction (Karoly et al. 2017) and that even multi-day rhythms are correlated to seizure occurrence (Baud et al. 2018).

In addition to examining the EEG signal properties originating from one or a few channels of interest, the use of functional connectivity patterns has been also suggested as a promising approach (Kramer et al. 2011; Lehnertz et al. 2014; van Mierlo et al. 2014; Geier et al. 2015a), as it may capture the underlying mechanisms in more detail and consequently improve our understanding of the emergence of epileptogenesis and ictogenesis. This is in agreement with recent evidence that has suggested seizure onset within a network of brain regions, challenging the traditional definitions of focal and generalized seizures (Lehnertz et al. 2014). Functional brain networks are often described using concepts from complex systems and network theory (Rubinov and Sporns 2010), aiming to quantify the interplay between the dynamic properties of network constituents (i.e., nodes and links) and the network topology. Similarly, to EEG signals, it has been shown that functional connectivity patterns are influenced by biological rhythms. For instance, the long-term properties of the functional brain networks of healthy subjects have been studied in (Ferri et al. 2007, 2008), where it was shown that these networks moved towards a small-world organization (high clustering coefficient and small characteristic path length) during the transition from wakefulness towards sleep. Furthermore, the same authors demonstrated that the small-world organization is slightly but significantly more evident over shorter time scales.

Global properties of epileptic networks around seizure onset have been characterized using measures such as clustering coefficient, shortest path length/ efficiency or synchronizability (Lehnertz et al. 2014). Additional studies have explored the relevance of local network properties, such as the importance of individual nodes (Koschützki et al. 2005; Rubinov and Sporns 2010), in the context of seizure dynamics (Kramer et al. 2008; Wilke et al. 2011; Varotto et al. 2012; Burns et al. 2014; Zubler et al. 2014; Geier et al. 2015a). Overall, studies related to seizure brain networks have suggested a transition from a more random functional network topology before seizure to a more regular topology during seizure, followed by a return to random topology after seizure, which may suggest a common mechanism of ictogenesis (Lehnertz et al. 2014). On the other hand, findings related to node-specific epileptic network characteristics have been less consistent, with important nodes not necessarily confined to the epileptic focus (Geier et al. 2013; Lehnertz et al. 2014; Geier and Lehnertz 2017).

The aforementioned changes in short-term network characteristics around seizure onset are contrasted by pronounced fluctuations of local and global network properties observed for the temporal evolution of epileptic brain networks over longer term periods (Kuhnert et al. 2010; Kramer et al. 2011; Geier et al. 2013). Specifically, the existence of long–term periodic fluctuations in the properties of functional brain networks was demonstrated in (Kuhnert et al. 2010), where it was also observed that these fluctuations exhibit larger amplitude compared to fluctuations due to seizure activity and status epilepticus. Averaging the normalized power spectral density estimates from all patients revealed a strong circadian component at around 24 h, with additional peaks observed at 12 and 8 h (Kuhnert et al. 2010). Geier et al. (2015b) investigated the long-term evolution of degree-degree correlations (assortativity) in functional brain networks obtained from patients with epilepsy. They observed large fluctuations in time-resolved degree-degree correlations which exhibited a periodic temporal structure which could be attributed to daily rhythms. Changes due to epileptic seizure onset, particularly possible pre-seizure alterations, were found to contribute marginally to the observed long-term fluctuations. (Geier et al. 2013) and (Geier and Lehnertz 2017) also showed that the functional brain networks of patients with epilepsy exhibit long-term fluctuations and that the epileptic focus is not consistently the most important node in the network, but node importance may vary drastically over time. Schelter et al. (2006) investigated the distribution of false seizure predictions during the day and their relation to the sleep-wake cycle, and revealed that the majority of false predictions occurred during non-REM sleep. Finally, Schelter et al. (2011) presented a strategy to avoid false seizure predictions by taking into consideration the influence of circadian rhythms and using adaptive thresholds.

Therefore, there is growing evidence that long-term periodic variations (hours/days) are present in both scalp/intracranial EEG signals and the corresponding functional brain connectivity patterns and that these periodicities possibly bear a relation to the occurrence of seizures. However, most epilepsy-related studies utilize short segments of data around seizures and attempt to detect short-term changes immediately preceding or following a seizure, assuming a relatively constant inter-ictal baseline. Given the long-term variations of this baseline, epileptic seizures and their detection/prediction may depend on the state of these variations. In turn, precise quantification of the relation between long-term periodic patterns in brain activity and seizure onset can only be performed using continuous, long-duration data obtained from patients with epilepsy.

In the present study, we rigorously investigate the long-term periodic properties of functional brain networks and their relation to seizure onset using long-duration scalp EEG data recorded from patients with epilepsy. Specifically, we construct functional brain networks and systematically examine their temporal structure over multiple time scales by using graph-theoretic measures to summarize network properties. The results reveal periodicities over different time scales both in summative network properties (average degree and efficiency) and network topology, whereby we use a novel measure based on the graph edit distance measure. Beyond the 24-hour circadian periodicity, we show that shorter periodicities at around 3 and 5 hours are consistently observed across all patients. The circadian and shorter periodicities result in network property fluctuations of higher amplitude compared to the fluctuations related to seizure onset *per se*. Furthermore, we address the question of whether seizures occur preferentially at specific phases of these periodic components. Using circular statistics, we show that seizure onset occurs preferentially at specific phases of these components. While also present for the circadian 24-hour periodicity, this relation is found to be stronger for the shorter periodicities (around 3 and 5 hours). This suggests that quantifying the characteristics of long-term network periodic fluctuations and their phase in particular may facilitate reliable detection of seizure onset. Finally, we show that correlations to seizure onset are much stronger for measures related from functional networks compared to periodicities in the EEG signals *per se*. Overall, our findings demonstrate the important role of biological rhythms on long-term functional connectivity patterns and seizure onset, which could be used for designing more reliable seizure detection and prediction algorithms.

## 2. Methods

### 2.1 EEG recording and preprocessing

Long-term video-EEG recordings were collected from 9 patients with epilepsy and one patient with psychogenic seizures in the Neurology Ward of the Cyprus Institute of Neurology and Genetics. The study was approved by the Cyprus National Bioethics Committee. All subjects gave written informed consent in accordance with the Declaration of Helsinki. Six patients were monitored using an XLTek (Natus Medical Incorporated, CA, USA) scalp EEG recording system (Patients 1-6), while the remaining four were monitored with a Nicolet (Natus Medical Incorporated, CA, USA) system (Patients 7-10). Table 9 summarizes the duration of the recordings, as well as the number and type of seizures of each patient. Seizures and sleep intervals were identified and marked by specialized neurophysiologists (coauthors ESP and SSP).

**Table 1:** EEG recordings

| Patient | Length of Recordings | Number of Seizures | Type of Seizures |
| --- | --- | --- | --- |
| 1 | 46h | 1 | Focal |
| 2 | 22h | 2 | Focal |
| 3 | 68h | 2 | Focal |
| 4 | 94h | 1 | Generalized |
| 5 | 36h | 1 | Generalized |
| 6 | 24h | 0 | Psychogenic |
| 7 | 21h | 1 | Focal |
| 8 | 71h | 2 | Focal |
| 9 | 27h | 6 | Generalized |
| 10 | 69h | 4 | Focal |

Twenty-one electrodes were placed according to the 10-20 international system with two additional anterotemporal electrodes. In addition, four electrodes were used to record the electrooculogram (EOG) and electrocardiogram (ECG) signals respectively. The data were recorded at a sampling rate of 200Hz and 500Hz for the XLTek and Nicolet systems respectively, using a cephalic reference, Cz, that was not part of the scalp derivations used to display the recorded channels. The EEG and EOG signals were band-pass filtered between 1 and 45Hz to remove line noise and muscle artifacts. The Lagged Auto-Mutual Information Clustering (LAMIC) algorithm (Nicolaou and Nasuto 2007; Daly et al. 2013) was applied to remove ocular artifacts using simultaneously recorded EOG recordings (2 channels) as reference signals (Ziehe and Müller 1998; Nicolaou and Nasuto 2003).

Subsequently, the data was converted to the bipolar montage, as this montage was found to be more robust to volume conduction effects in the present case, where a limited number of electrodes was available (Christodoulakis et al. 2013). According to this montage, pairs of EEG electrodes placed in nearby locations of the scalp are used to obtain the time-series by subtracting the corresponding measurements, forming the pairs Fp1-F7, F7-T3, T3-T5, T5-O1, Fp2-F8, F8-T4, T4-T6, T6-O2, Fp1-F3, F3-C3, C3-P3, P3-O1, Fp2-F4, F4-C4, C4-P4, P4-O2, Fz-Cz and Cz-Pz. In general, the main results did not change when different montages were used.

### 2.2 Functional brain network construction

Each bipolar time series, e.g. Fp1-F7, corresponds to a node on the network, representing the area on the scalp that lies between the two electrodes, i.e. the area between Fp1 and F7 (Figure 1). Note that nodes do not change over time. We identified edges (i.e. connections) between nodes by quantifying time- and frequency-domain correlations between the corresponding EEG time series. To track network related changes over time, we used 5- second non-overlapping windows. Specifically, for each window, we quantified the correlation between all time-series pairs, using one of the following measures: cross-correlation, corrected cross correlation, and coherence (subsections 2.2.1-2.2.3). Finally, we constructed binary graphs by using a threshold that is specific to each correlation measure as discussed in Section 2.2.4.

#### 2.2.1 Cross-correlation

For any pair of time series, *x*(*t*) and *x*(*y*), the normalized cross-correlation is calculated as follows:

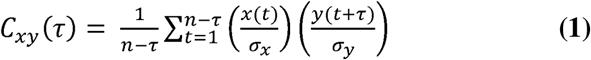

where σ _*x*_ and σ _*y*_ are the standard deviations of *x* and *y* respectively. *C*_*xy*_ is computed for a range of values for the lag *τ*, which depends on the sampling frequency; for the required range of [-100 100] ms chosen here, *τ* lies between [-20 20] and [-50 50] time lags for the data recorded with the XLTek and Nicolet systems respectively. C_*xy*_ takes values between −1 and 1, with 1 indicating perfect linear positive correlation, −1 perfect linear negative correlation and 0 no correlation. The maximum of the absolute value of cross-correlation, *max*_*τ*_| *C*_*xy*_|, over the chosen range of values, is used to quantify the degree of correlation between the two signals within a given window.

#### 2.2.2. Corrected cross-correlation

Cross-correlation often takes its maximum at zero lag in the case of scalp EEG measurements. Consistent zero-lag correlations could be due to volume conduction effects, whereby currents from underlying sources are conducted instantaneously through the head volume to the EEG sensors (i.e., assuming that scalp potentials have no delays compared to their underlying sources (quasi-static approximation) (Nunez and Srinivasan 2006; Christodoulakis et al. 2013). In principle, true direct interactions between any two physiological sources will typically incur a nonzero delay due to transmission speed, provided that the sampling frequency is high enough to capture such delays. Therefore, to measure true interactions not occurring at zero lag, we calculated the corrected cross-correlation, which is a measure of the time-series’ asymmetry, as defined in (Nevado et al. 2012), by subtracting the negative-lag part of *C*_*xy*_(*τ*) from its positive-lag counterpart:

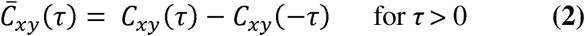

Note that 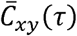 provides a lower bound estimate of the nonzero-lag cross correlations and is typically much smaller than *C*_*xy*_. As in the case of cross-correlation, the maximum within the selected range of time lags ([-100 100] ms) was used to quantify correlation.

#### 2.2.3 Coherence

Coherency may be viewed as a measure of cross-correlation in the frequency domain; it measures the linear correlation between two signals *x* and *y* as a function of the frequency *f*. It is defined as the cross-spectral density *S*_*xy*_(*f*), between *x* and *y* normalized by the auto-spectral densities, *S*_*xy*_(*f*) and *S*_*xy*_(*f*), of and respectively. Coherency is a complex number, as the cross-spectral density is complex, whereas the auto-spectral density functions are real. Therefore, in many cases coherence (or the squared coherence), which is defined as the magnitude of coherency (or its square), is employed as a measure of correlation in the frequency domain, i.e.:

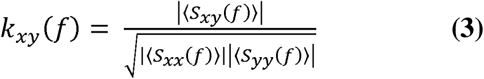

The value of *k*_*xy*_(*f*) ranges between 0 and 1, with 1 indicating perfect linear correlation and 0 no correlation between *x* and *y* at frequency *f*. We calculated the maximum coherence value between EEG signals for all nodes both for the broadband signals (1-45Hz), as well as within the following frequency bands; delta (1-4Hz), theta (4-8Hz), alpha (8-13Hz), beta (13-30Hz) and gamma (30-45Hz). The maximum coherence value within each frequency band was used to quantify correlation.

#### 2.2.4 Network binarization

To obtain a binary (rather than weighted) network, we used a threshold to define strongly correlated nodes, which were subsequently assigned edges with a weight of 1. The threshold value was selected such that networks with similar average degrees were obtained for different correlation measures. For the three examined correlation measures (as well as alternative ones - see (Christodoulakis et al. 2013), it was found that different threshold values yielded very similar results in terms of the time evolution and the corresponding periodicity patterns for all network measures. This was found to be the case except when the threshold value was too high (e.g. close to one for correlation or coherence) yielding disconnected graphs, or too low (close to zero), yielding densely/fully connected graphs. For threshold values between these two extremes the graphs exhibited similar properties, as shown in (Christodoulakis et al. 2013) for shorter data segments, and in supplementary material (Fig. A.1) for a representative long-duration data set (94 hours for Patient 4). From Fig. A.1 it is evident that the observed patterns and periodicities of the network properties (average degree in this figure) are similar for threshold values between 0.2 and 0.8. As the temporal patterns of the functional brain network measures – as opposed to their absolute values – are of interest here, the precise threshold value was not a confounding factor. Hence, we have chosen the threshold independently for each correlation measure, aiming to obtain a similar average degree values for all measures. Specifically, for cross-correlation and coherence, the threshold was set to 0.65, while for corrected cross-correlation it was set to 0.20. We also used surrogate data to construct binary networks using the original time series by randomizing their phase (Theiler et al. 1992); however, they were found to yield rather densely connected networks and the results were similar to those obtained by simply setting a fixed, low threshold value. Therefore, we present results using the simple thresholding method.

### 2.3 Functional brain network evolution

For each subject, the evolution of functional brain networks over time was quantified in two ways. First, we quantified how different properties of the graph changed over time, namely average degree, global efficiency and clustering coefficient (subsections 2.3.1-2.3.3). Furthermore, we also performed direct comparisons of the network topologies at different times by means of the graph edit distance, which quantifies the dissimilarity between different graphs (Section 2.3.4). In the following, let *n* denote the number of nodes of the network (in our case *n*=18) and *N* the set of all nodes.

#### 2.3.1 Average degree

The degree *k*_*i*_ of a node *i* is defined as the number of nodes *j* in the network to which node *i* is connected to via an edge, *e*_*ij*_; that is, the number of edges incident to *i*. The average network degree is given by:

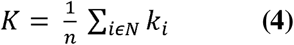

The average degree of a graph quantifies how well connected the graph is.

#### 2.3.2 Global efficiency

Although the average degree reflects the average number of connections any network node may have, it does not provide any information regarding the actual distribution of the edges and, hence, how easy it is for the information to flow in the network. This can be captured by the shortest (or geodesic) path length *d*_*ij*_ between a pair of nodes *i* and *j*. It is defined as the minimum number of edges that have to be traversed to get from  node *i* to *j*. The characteristic path length is defined as the average shortest path length over all pairs of nodes in the network:

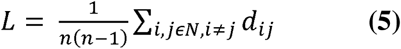

However, the characteristic path length is well defined only for pairs of nodes that are connected through a path. If any two nodes *i* and *j* are not connected through a path, the shortest path length between them is *d*_*ij*_=∞, hence the average shortest path length for the network becomes *L* = ∞. A workaround for this is to consider only pairs of nodes that are connected through a path, but this does not reflect the connectivity of the entire network. To overcome this, Latora and Marchiori (2001) defined the efficiency between a pair of nodes as the inverse of the shortest distance between the nodes, 1/*d*_*ij*_. The global network efficiency is subsequently defined as the average efficiency over all pairs of nodes (Latora and Marchiori 2001):

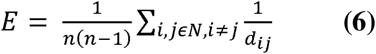

Therefore, when a path between two nodes does not exist, the corresponding efficiency is zero.

#### 2.3.3 Clustering coefficient

A cluster in a graph is a group of nodes that are highly interconnected. The clustering coefficient (Watts and Strogatz 1998) **C**_*i*_ of a node *i* is defined as the fraction of existing edges between nodes adjacent to node *i* over the maximum possible number of edges:

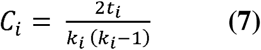

where *k*_*i*_ is the degree of node *i,* and *t*_*i*_ denotes the number of edges between *e*_*jj′*_ pairs of nodes *j* and *j′* that are both connected to *i*. The clustering coefficient, *C*, of the network is the mean clustering coefficient across all nodes.

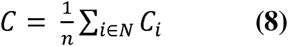

#### 2.3.4 Graph edit distance

Using the graph properties defined above, the general characteristics of two (or more) functional brain networks corresponding to different times may be compared. If the value of a network property (e.g. degree, clustering coefficient, efficiency) corresponding to one network differs substantially from the value of the same property corresponding to some other network, it is reasonable to assume that these two networks differ in their topology as well. However, comparing networks with similar network summative measure values can be inconclusive. Consider, for instance, two networks with average degree 5. It is impossible to conclude whether these two networks are similar in structure by the average degree alone; for example, one of the networks could have its edges equally distributed among the nodes (each node having degree 5), while the other could have a large number of low degree nodes and a few hubs possessing a large number of connections. In order to more accurately quantify topological difference between graphs, we compared every pair of graphs directly in terms of their structure, using the graph edit distance measure. The graph edit distance (Dickinson et al. 2003) estimates graph similarity in terms of the minimum number of insertions and deletions of either nodes or edges that must take place in one graph in order to make it identical to the second. When the two graphs share the same set of nodes, as is the case for EEG functional brain networks, the distance between the graphs is defined as the minimum number of edge insertions and deletions that are necessary in order to make the two graphs identical. Essentially, the graph edit distance *g*_*AB*_ between two functional brain networks *A* and *B* is equal to the number of edges that exist only in one of the two graphs. Note that graph edit distance is a symmetric measure, since an insertion of an edge in one graph is equivalent to a deletion in the other.

#### 2.3.4 Periodicity estimation

To characterize the periodicities that arise in functional brain network characteristics over time, we investigated the evolution of both the resulting graph-theoretical summative network properties and the network structure. Each of the three summative network properties - average degree, global efficiency, and clustering coefficient - provides a single value per network, yielding a single time series per measure. To characterize the periodic structure of these time series, we used the Lomb-Scargle (LS) periodogram to estimate the power spectral density (PSD) (more details are given in the Supplementary material – Section A.2).

As mentioned earlier, similarity between network properties does not necessarily imply similarity in network structure; consequently, periodicity in network properties does not necessarily imply structure/topology periodicity. For this reason, we investigated the existence of periodicities in the network structure by developing a novel measure based on the graph edit distance. Specifically, we consider the course of the functional brain network over time as a vector, ***A***, of graphs, whereby ***A****(t)* corresponds to the functional brain network at time lag *τ* (and each time lag corresponds to one 5-sec window). We then compare this vector with shifted copies of itself and for each shift value *τ*, we calculate the average graph edit distance between all network pairs that are separated by *τ* time lags:

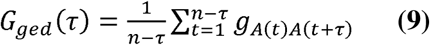

Similarly, to the autocorrelation function, this measure can reveal periodic changes in the network structure over time, complementing network summative properties. Note that, for any shift value *τ*, the lower the average graph edit distance value *G*_*ged*_(*τ*) is, the more similar the corresponding graphs are.

#### 2.3.5 Circular Statistics

To investigate the relation of seizure onset to network periodicities, we calculated the instantaneous phase at seizure onsets for each of the main identified periodic components (Section 3.2) and obtained a phase distribution for each of these components. Subsequently, we used circular statistics to examine whether seizure onsets occurred at specific/preferred phases (as opposed to random phases). To this end, we first performed zero-phase digital filtering of the average degree time series to obtain band limited signals around the main identified periodicities (±0.5h before and after the main period of each component on a subject-to-subject basis). Specifically, we investigated the periodic components at 3.6, 5.4, 12 and 24 h (mean values across subjects), as these were consistently found for all patients (Table 3). Similar periodic components were obtained using the other two examined connectivity measures (clustering coefficient and efficiency) for all patients, but here we present the results only for average degree. Note that, as mentioned above, subject-specific bandlimited time-series were obtained around the main period identified for each subject to account for physiologically expected individual differences. Subsequently, we applied the Hilbert transform (Klingspor 2015) to the resulting bandlimited signals in order to calculate the instantaneous phase of each periodic component at the time of seizure onset for all patients and seizures. The same procedure was followed for obtaining the EEG signal power to investigate long-term periodic components exist and their correlation to seizure occurrence. Specifically, we used the averaged broadband (1-45Hz) EEG signal power for identifying the main periodic components. The resulting final phase distributions for each periodic component were subsequently investigated using circular statistics.

Circular statistics are suitable for data that are defined within an angular scale, as in the present case, whereby there is no designated zero and, in contrast to a linear scale, the designation of high and low values is arbitrary (Berens 2009). In this context, the mean resultant vector after transforming the data points to unit vectors in the two-dimensional angular plane is given by:

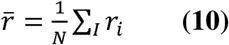

where **r**_*i*_ is the unit vector. The length of the mean resultant vector is a crucial quantity for the measurement of circular spread or hypothesis testing in directional statistics. The closer it is to one, the more concentrated the data sample is around the mean direction. The resultant vector length is computed by:

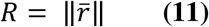

The circular variance, which is related to the length of the mean resultant vector (Berens 2009), is defined as:

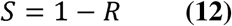

In contrast to the variance on a linear scale, the circular variance *S* is bounded within the interval [0, 1]. It is indicative of the spread in a data set. If all samples point towards the same direction, the corresponding mean vector will have length close to 1 and the circular variance will be small. If the samples are spread out evenly around the circle, the mean vector will have length close to 0 and the circular variance will be close to its maximum value of one. We investigated whether phase values at seizure onset times were distributed uniformly around the circle from 0 to 2π or whether a common mean direction existed by using the instantaneous phase obtained for all seizures and patients. We applied the Rayleigh test to assess significance with the following common null hypothesis H_0_: the population is distributed uniformly around the circle (Fisher 1993). The Rayleigh test is particularly suited for detecting a unimodal deviation from uniformity. The approximate *p*-value under H_0_ is computed as (Zar 1999):

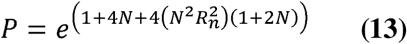

where *Rn* = *R* × *N* where *N* is the number of observations. A small *p* value indicates a significant departure from uniformity and indicates rejection of the null hypothesis. This was done for all seizures (twenty in total) corresponding to nine different patients.

To account for the fact that we had multiple seizures for some subjects, we also created groups of nine samples (one seizure per patient) for all possible combinations and investigated periods (3.6h, 5.4h,12h and 24h) in order to perform correction for multiple comparisons (Zar 1999). Note that for the 24 h circadian periodic component, these groups included six samples only, since the recordings of only six patients were longer than 24 hours. The corrected *p*-values were computed by applying the Rayleigh test (Zar 1999). All the above quantities were computed with the Matlab (Mathworks, Natick MA) CircStat toolbox (Berens, 2009).

## 3 Results

The individual results presented below correspond to Patient 4, who yielded the longest recording (94 hours). Similar results were obtained from all ten patients, as discussed in Subsection 3.6.

### 3.1 Periodicities in network properties

#### 3.1.1 Time domain

Figure 1 (a-c) shows the time course of the three summative network properties of interest (average degree, global efficiency and clustering coefficient) in the case of standard cross-correlation. It can be seen that the obtained functional brain networks were less connected and less clustered when the patient was awake compared to sleep (grey shaded bars). This pattern occurs periodically, in cycles of approximately 24 hours, a fact that is further illustrated by the corresponding autocorrelation sequences (Figure 1 (e-f)), which also demonstrate a clear periodic pattern with a main period equal to around 24 hours. Very similar results were obtained when the corrected cross-correlation was used for constructing the networks (Figure 2). In addition to the main 24-hour cycles, it can be observed that additional periodic components at smaller time scales co-exist in the time course of the obtained functional brain network measures; note, for example, the spikes that occur at both awake- and sleep-times separated by approximately 75 minutes. These weaker periodicities are examined in detail in Subsection 3.2. In the supplementary material, we show the 99% confidence interval of the difference between the network average degree during sleep time minus the average degree during awake time for all patients (Figure A.2).

**Figure 1:**
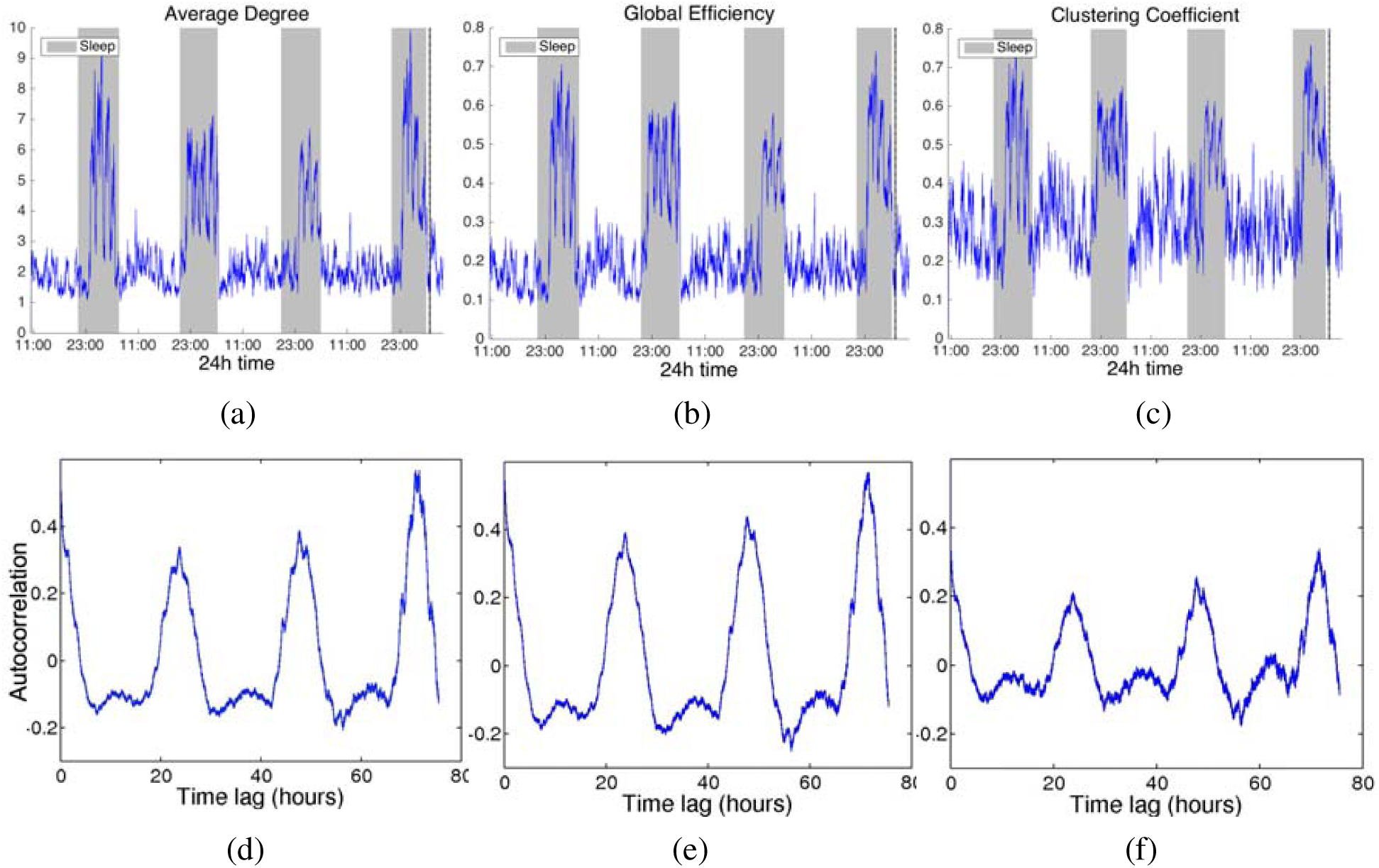
Top: Network average degree (a), global efficiency (b), and clustering coefficient (c) for Patient 4 as a function of time, obtained using cross correlation for quantifying pairwise correlations. For presentation purposes, the obtained network properties have been smoothed using a moving average filter. The vertical dashed line indicates seizure onset and the grey bars indicate sleep intervals. Bottom row: Corresponding autocorrelation sequences. A periodic pattern with a main period equal to around 24 hours can be observed. Functional brain networks during sleep periods were found to be more connected and clustered compared to awake periods.

**Figure 2:**
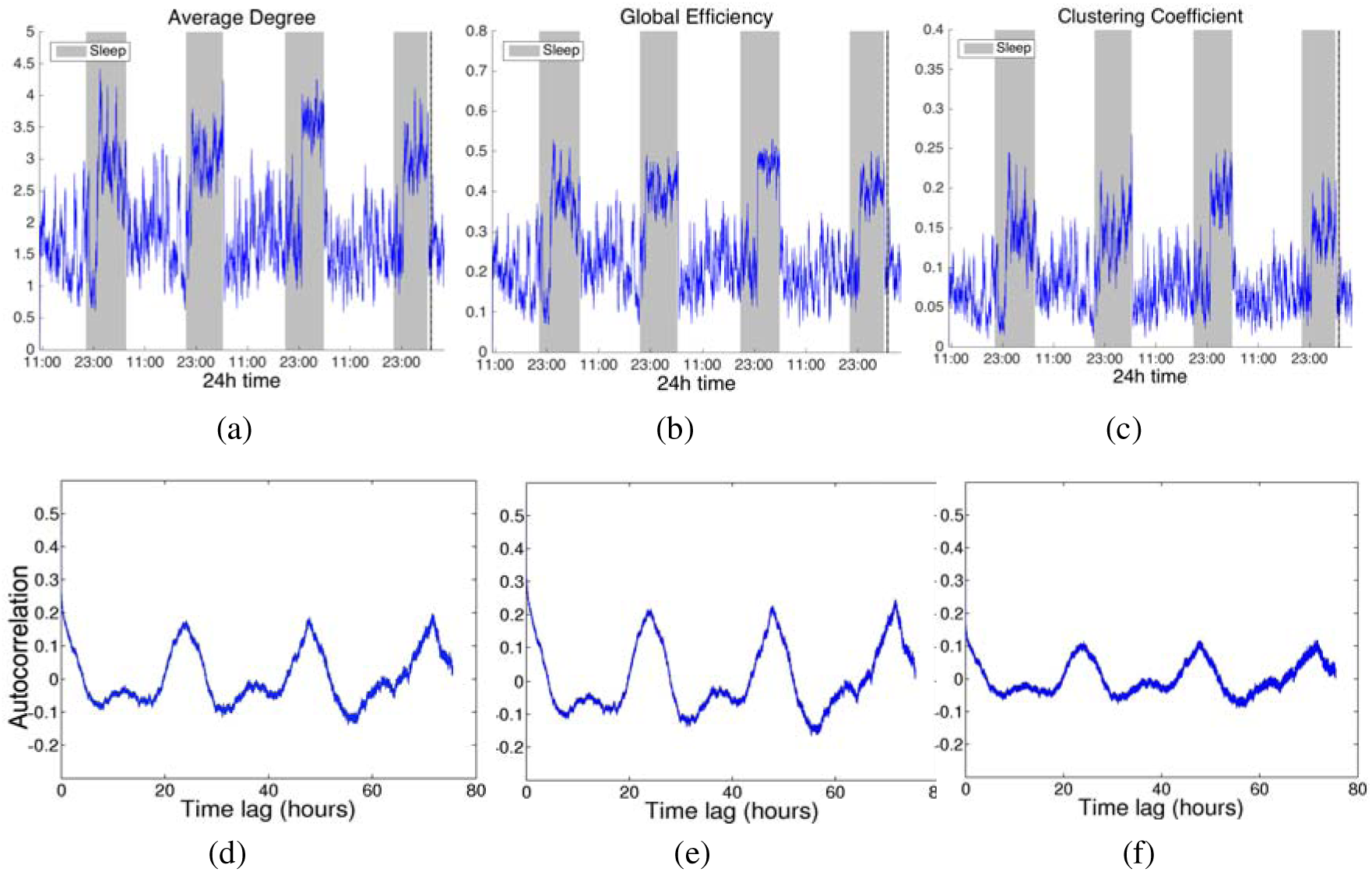
Network average degree (a), global efficiency (b), and clustering coefficient (c), and their corresponding autocorrelation sequences, respectively, (d)-(f) for Patient 4 as a function of time obtained using corrected cross correlation for quantifying pairwise correlations. For presentation purposes, the obtained network properties have been smoothed. The vertical dashed line indicates seizure onset and the grey bars indicate sleep intervals. The observed patterns are very similar to those obtained when using standard cross correlation (Fig. 2).

#### 3.1.2 Frequency domain

The network average degree obtained when using coherence within the six frequency bands of interest is shown in the supplementary material (Figure A.2). Global efficiency and clustering coefficient yielded similar periodicities within all bands and are not shown separately. Except for the gamma band, the 24-hour periodicity is clear for all other frequency bands, particularly for the alpha band, followed by the beta band, and finally the delta and theta bands. Overall, these results suggest that the alpha and beta bands dominate the long-term broadband network properties.

### 3.2 Periodicities at shorter time scales

In addition to the main 24h periodicity observed for all network measures, periodicities at shorter time scales can also be observed (Figs. 1-2, A.2). In this section, we investigate these periodicities in more detail. Figure 3 shows the LS periodogram for the average degree of Patient 4 obtained using cross-correlation and corrected cross-correlation (Figs. 1 and 2 respectively). The peaks in the periodogram correspond to different periodic components and have been marked accordingly.

Table 2 summarizes the main periodic peaks consistently identified for all subjects, whereby a peak was considered consistent if its location differed by half an hour at most across subjects. Note that the 24h periodicity is absent for patients whose recordings span less than 24h. Periodic components were consistently observed at the following mean locations: 3.4 h, 5.6 h, 12h and 24h, while component with an average period of 1.7 h was identified in six out of 10 patients.

**Figure 3:**
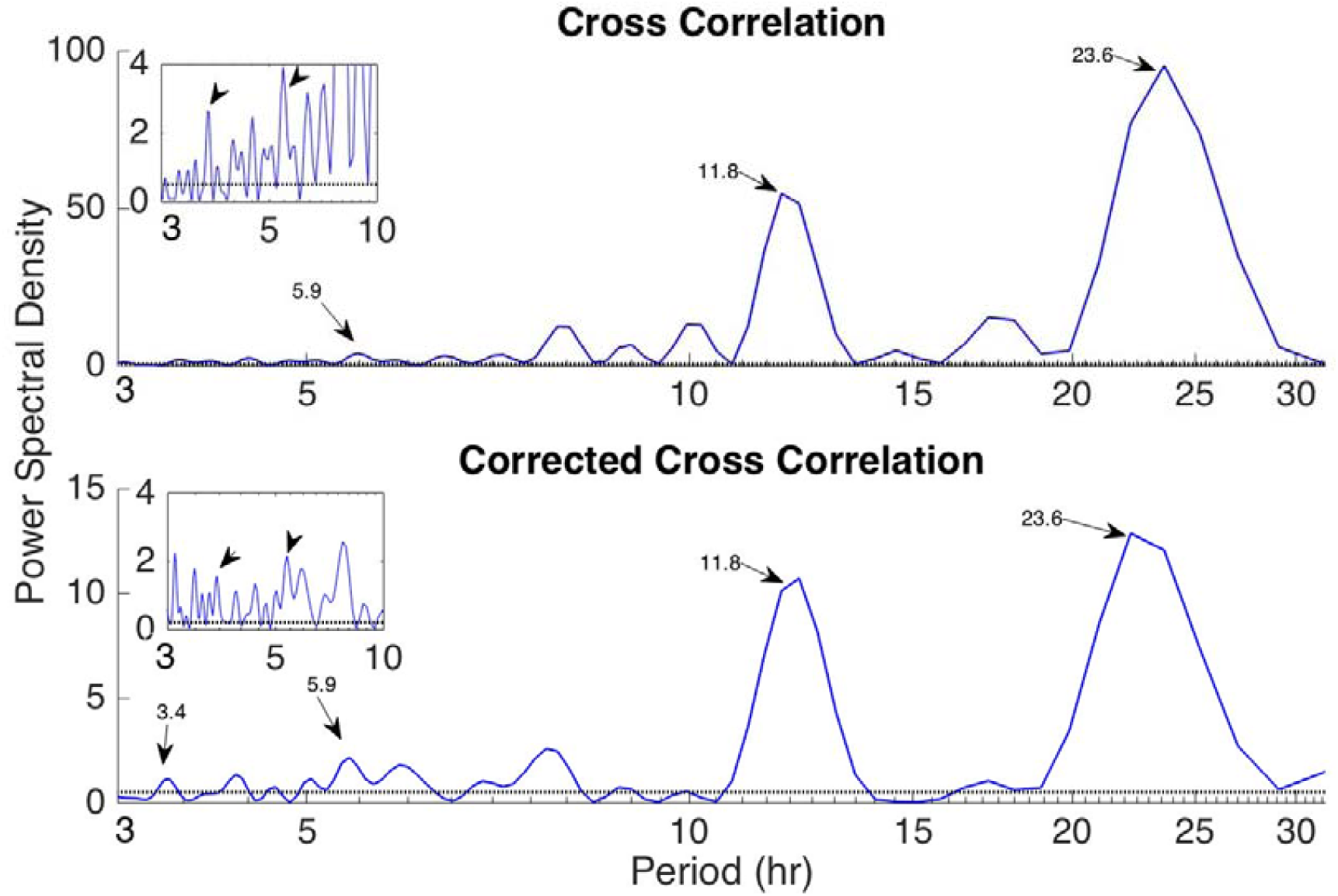
Periodogram of the time-resolved average degree of the functional brain networks of Patient 4 using cross-correlation (top panel) and corrected cross correlation (bottom panel). The inset graphs show a zoomed-in version for periods between 3-10 h. The periodic components that were consistently identified for all subjects (located at 3.4 h, 5.9 h, 11.8 h and 23.6 h for this particular subject) are marked on the plot. The dotted horizontal lines denote the statistical significance level (*p*=0.05).

**Table 2:** Main periodic components identified in the network average degree for all subjects.

| Periodic peak<br>location mean<br>(range) | P1 | P2 | P3 | P4 | P5 | P6 | P7 | P8 | P9 | P10 |
| --- | --- | --- | --- | --- | --- | --- | --- | --- | --- | --- |
| 24.0 (23.6-24.5) | √ | N/A | √ | √ | √ | N/A | √ | √ | N/A | √ |
| 12.0 (11.8-12.2) | √ | √ | √ | √ |  | √ |  | √ | √ | √ |
| 5.4 (4.8-5.9) | √ | √ | √ | √ | √ | √ | √ | √ | √ | √ |
| 3.6 (3.2-3.8) | √ | √ | √ | √ | √ | √ | √ | √ | √ | √ |
| 1.7 (1.7-1.9) | √ | √ |  | √ |  | √ |  | √ | √ |  |

### 3.3 Periodicities in network topology

In addition to examining the evolution of properties of functional brain networks over time, we also investigated evaluated the evolution of their topology using the average graph edit distance (Equation (9), Section 2.4). Figure 4 shows the results for Patient 4, while results for all patients are presented in the supplementary material (Figure A.5). In this case, note that similarity between graphs at different time points is now indicated by the presence of a local minimum. The main periodic peaks revealed but the LS periodogram of the average graph edit distance (Fig. 4, bottom panel) are in agreement to those identified using the average degree (Figs. 1-2). In a similar fashion, the periods identified for the remaining nine patients were similar to those identified by their summative network properties, suggesting that network topology is characterized by the same periodic components.

**Figure 4:**
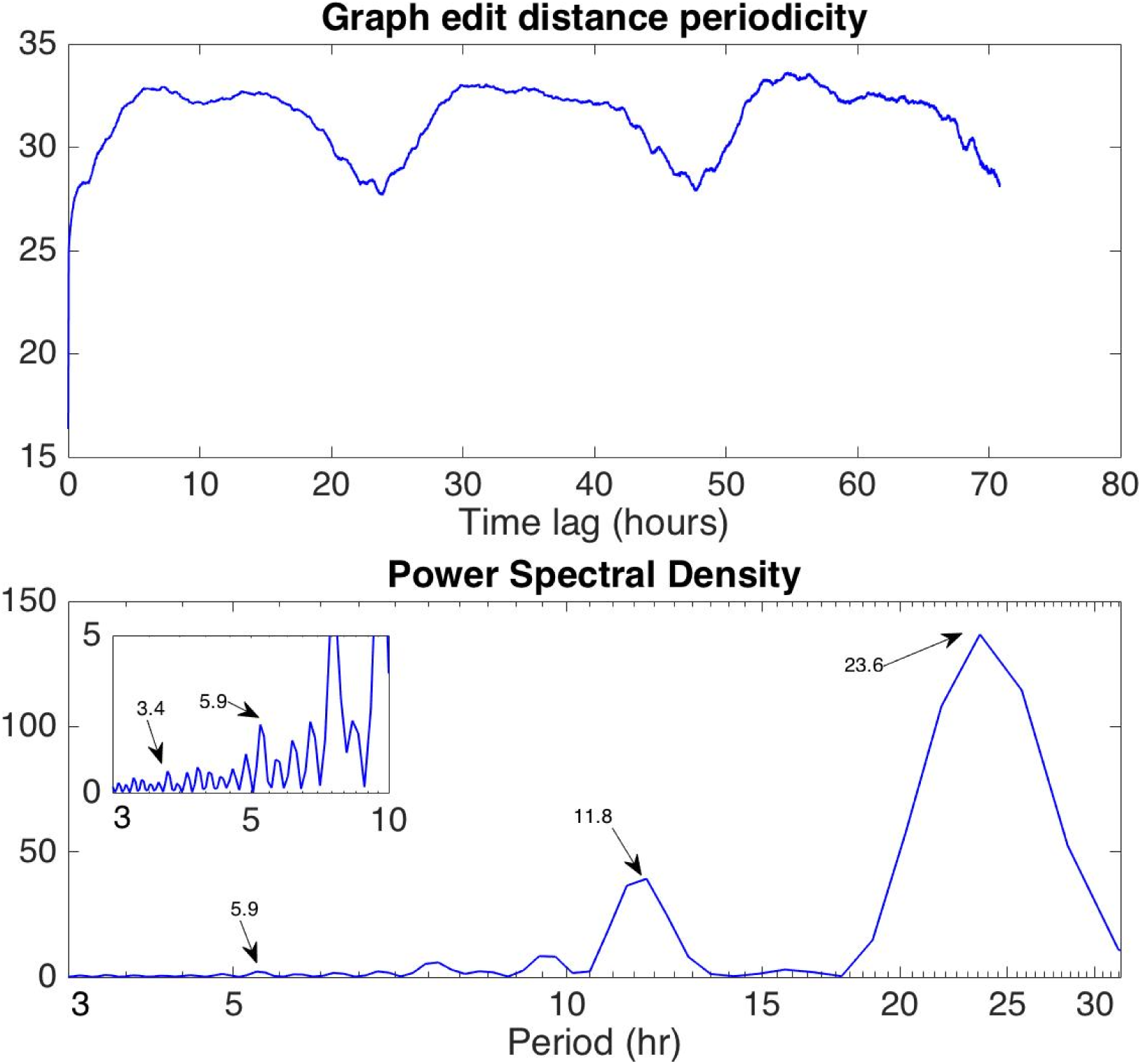
Periodicity in the network structure as assessed with the average graph edit distance for Patient 4 (top panel). The PSD of the average graph edit distance indicates periodic components at 3.4, 5.9, 11.8 and 23.6 h (bottom panel) in agreement to the average degree (Fig. 1-2). The inset shows the zoomed in PSD for periods between 3-10 h.

### 3.4 Periodicities in EEG signal power

To complement the results on the evolution of functional brain networks over time, we additionally investigated the periodicities in the recorded EEG signals. Specifically, we calculated the power of the EEG signals averaged over all channels within the six frequency bands of interest (broadband, delta, theta, alpha, beta and gamma) and show the obtained results in the Supplementary material for Patient 4 (Figure A.5). Results were found to be overall similar across subjects. The main circadian periodicity is evident mostly in the broadband, beta and gamma signal power. Compared to the observed periodicities in network summative properties (average degree; Fig. A.3), there exist similarities (especially for the beta band but also for the theta and delta bands). However, pronounced differences can be observed for the alpha and gamma bands; in the former case, only the average degree yielded a clear circadian periodicity (Fig. A.3(d)), while in the latter case this was found to be the case for the EEG signal power only (Fig. A.5(f)). Finally, the results suggest that during sleep, the EEG delta and theta power increased (Fig. A.5(b-c)), while beta and gamma power decreased (Fig. A.5(e-f)) which suggests an overall slowing of the sleep EEG in agreement to previous studies (Bazil and Walczak 1997; Minecan et al. 2002; Bruzzo et al. 2008).

### 3.5 Effects of seizure onset on connectivity measures

Figure 5 shows the average degree of Patient 4 obtained using cross-correlation (Fig. 1(a)) within intervals of increasing duration around the seizure, which occurs at approximately 91 hours after the beginning of the recording. In Figs. 5 (a-b), where ±2 and ±5 minutes around the seizure onset are plotted, an increase in the average network connectivity slightly before seizure onset can be observed, which gradually decreases after the onset until it reaches a lower level compared to the pre-seizure connectivity. However, when longer intervals are considered (Figs. 5 (c-f), where intervals of ±15 minutes or longer are shown), it can be observed that the changes in network connectivity occurring very close to seizure onset have a small amplitude compared to the slower fluctuations that are inherent in network connectivity (e.g. those occurring during the transition from sleep to awake and vice versa). This was a consistent observation across different seizures and subjects and implies that it may be difficult to distinguish seizure-related from physiologically-related connectivity changes.

**Figure 5:**
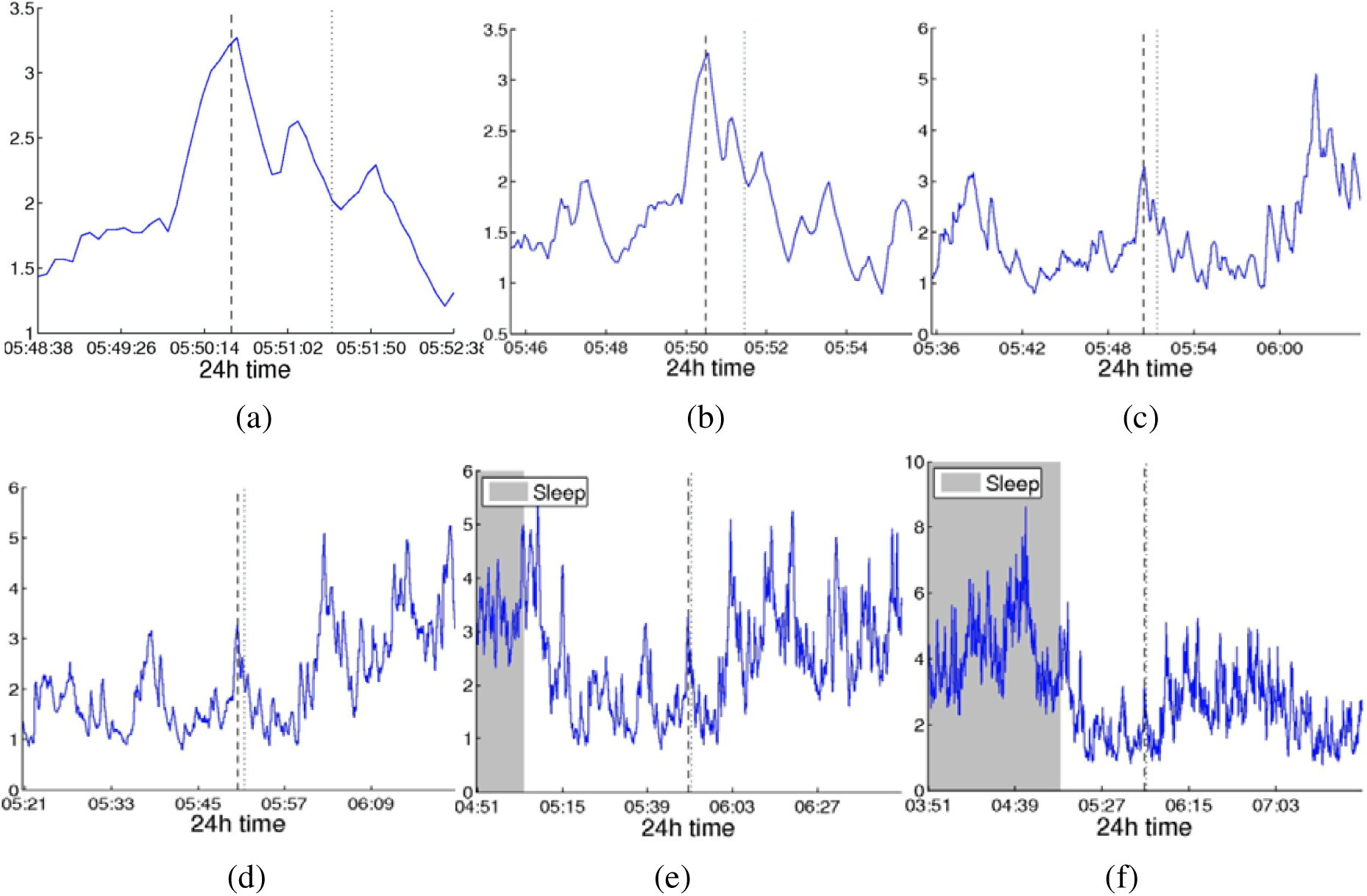
Average degree obtained using cross-correlation around seizure onset for time intervals of increasing duration (Patient 4): (a) ±2 minutes around seizure onset, (b) ±5 min, (c) ±15 min, (d) ±30 min, (e) ±1 hour, (f) ±2 hours. Seizure onset and end are indicated by the dashed and dotted vertical lines respectively. Whereas for smaller intervals (±2 to ±5 minutes – panels a, b) an increase in the average degree just before seizure onset, followed by a decrease persisting after seizure end, can be observed, the amplitude of this change is considerably smaller when compared to longer term fluctuations in network connectivity (panels c-f).

### 3.6 Correlation of functional brain network periodicities to seizure onset

The instantaneous phases of the main identified periodicities (mean periodicities: 3.6h, 5.4h, 12h and 24h) both for the average network degree and the EEG signal power, are shown in Figure 6 for all seizures from nine patients. Note that no seizures were recorded for Patient 6; therefore, this patient has not been included in the present analysis. The left panels show the instantaneous phases on the unit circle and the right panels show the corresponding angular phase distribution histograms. The blue and green circles (left panels) denote the instantaneous phase at seizure onset of the average degree and EEG signal power respectively, for each periodic component and for all seizures and patients. The blue and green lines (right panels) indicate the direction and magnitude of the mean resultant vector R of the average degree and EEG signal power respectively.

A mean resultant vector R with a larger magnitude (close to one) suggests that the data sample is more concentrated is around the mean direction. Therefore, it can be observed that the average degree yields distributions that is more concentrated around their mean, particularly for the shorter (3.4h and 5.6h) periodic components. This is further illustrated in Table 3, where the corresponding values of the mean resultant vector length R are given. The values obtained for the average degree suggest that the instantaneous phases are not distributed uniformly, but seizure onset occurs within specific phase ranges. In contrast, the instantaneous phases obtained through power are more uniformly distributed around the circle. Statistically, we examined the departure from uniformity using Rayleigh’s test after correcting for the presence of multiple seizures in some subjects as described in Section 2.3.5. The resulting p-values are given in Table 4 and they suggest that the null hypothesis (uniform distribution) was rejected in all cases for the average degree but not for the EEG signal power. The largest departures from uniformity (smaller p-values) were obtained for the shorter periodicities. These results suggest that long-term brain connectivity is more strongly correlated to seizure onset compared to the EEG signal properties.

**Figure 6:**
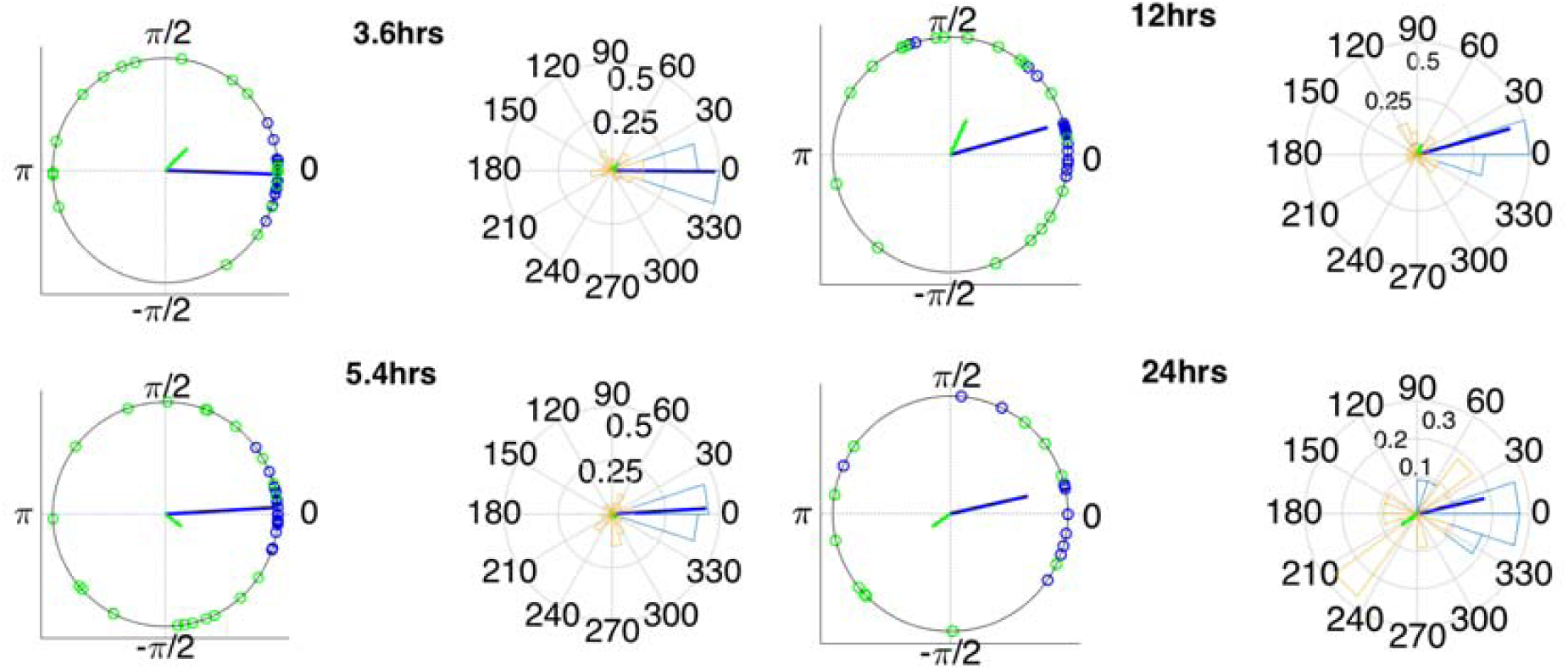
Instantaneous phases of the network average degree and EEG signal power at seizure onset for the main identified periodicities (3.4, 5.6, 12 and 24 h). Left panels: location of instantaneous phases at seizure onset on the unit circle. Right panels: Angular histograms of the corresponding phase distributions. The blue (green) circles denote the instantaneous phase for each periodic component of the average degree (EEG signal power) at seizure onset for all seizures and patients and the blue (green) lines denote the corresponding mean resultant vector (R). For all periodic components, particularly the 3.4 and 5.6 h components, the phase distribution is pronouncedly different from a uniform distribution, but only for the average degree.

**Table 3:** Mean resultant vector length (R) values for the phase distributions obtained from network average degree and EEG signal power. In all cases, the average network degree yields a vector that is more concentrated around its mean value, suggesting a clear correlation between seizure onset and instantaneous phase, particular for the 3.4 and 5.6 h components.

| Signal | R |  |  |  |
| --- | --- | --- | --- | --- |
|  | 3.6h | 5.4h | 12h | 24h |
| Average Degree | 0.97 | 0.98 | 0.85 | 0.72 |
| Power | 0.28 | 0.18 | 0.33 | 0.19 |

**Table 4:**
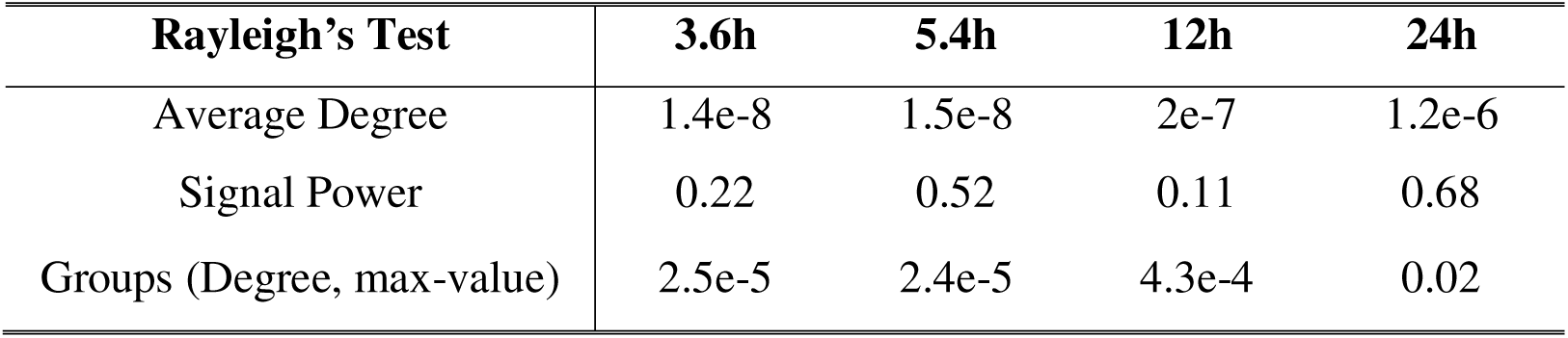
P-values corresponding to Rayleigh’s Test for comparing the instantaneous phase distributions to the uniform distribution. In all cases, the null hypothesis (uniform distribution) is rejected with high confidence for the average network degree, in contrast to the EEG signal power.

| Rayleigh’s Test | 3.6h | 5.4h | 12h | 24h |
| --- | --- | --- | --- | --- |
| Average Degree | 1.4e-8 | 1.5e-8 | 2e-7 | 1.2e-6 |
| Signal Power | 0.22 | 0.52 | 0.11 | 0.68 |
| Groups (Degree, max-value) | 2.5e-5 | 2.4e-5 | 4.3e-4 | 0.02 |

## 4 Discussion

We have examined the time evolution of functional brain networks in patients with epilepsy using long duration scalp EEG measurements (between 22 and 94 hours). These brain networks were constructed using three different correlation measures: cross-correlation, corrected cross-correlation and coherence. Their time evolution was monitored using three widely used summative network measures: average node degree, global efficiency and clustering coefficient. For all three correlation measures, it was found that the network properties exhibited a main 24 h periodicity, as well as additional periodicities at shorter time scales. Importantly, a strong correlation between these periodic components and seizure onset was revealed by examining the distribution of their instantaneous phases at seizure onset, particularly for the shorter periodicities. Furthermore, the same correlation was not observed between seizure onset and the periodic components of the EEG signal power. Overall, these findings suggest that functional network properties and specifically average connectivity are a more specific marker of the probability of seizure onset and that they could be taken into account for designing more robust seizure detection and prediction algorithms, which has proven over the years to be a notoriously difficult task, with reproducible results being hard to achieve. Our results suggest that one of the reasons is the fact that is often ignored that inter-ictally the EEG signals, and consequently the resulting functional brain networks, are far from being constant.

The long-term patterns (several hours to days) of EEG signal and functional brain network properties have been investigated; however, this has been done mainly using intracranial recordings (Kreuz et al. 2004; Schad et al. 2008; Kuhnert et al. 2010; Kramer et al. 2011; Schelter et al. 2011; Geier et al. 2013, 2015b; Geier and Lehnertz 2017). The use of scalp EEG has been considerably more limited, due to the practical difficulties of collecting long-term scalp EEG data. For instance, studies that have investigated the modulation of scalp EEG signals by circadian and ultradian rhythms have used short data segments collected at different times of the day (Aeschbach et al. 1997, 1999). Furthermore, studies focusing on the effect of sleep (Ferri et al. 2007, 2008) have used scalp EEG overnight recordings. In another study, (Schad et al. 2008) used both long-term intracranial and scalp EEG data from six patients to perform seizure detection and prediction by using integrate-and-fire neuron models to estimate spiking rates. A clear 24-h pattern was reported in the feature that quantified spike rate. Also, even though iEEG achieved overall better performance, scalp EEG performed better for some patients.

Several more recent studies have used functional connectivity in the context of seizure prediction and detection (Van Mierlo et al. 2011; van Mierlo et al. 2014). However, in most of these studies this was done using relatively short time windows around the seizure to perform seizure prediction and localization of the epileptogenic focus. In a more recent study (Campo et al. 2018) investigated the underlying brain network dynamics before seizure onset using intracranial video EEG data in 10 patients with pharmacoresistant epilepsy. Specifically, they constructed weighted undirected graphs using iEEG data obtained several hours before each seizure and the corresponding brain networks were characterized using eigenvector centrality. Using this approach, they showed that the preictal phase consists of two sequential events: a state characterized by networks that are more spatially homogeneous, with a (patient-specific duration) of 2-8 h, followed by a state of decreased connectivity with a duration of approximately 30 min before onset. Furthermore, several studies have used iEEG to examine the longer term properties of functional brain networks in patients with epilepsy and their relation to seizure onset (Kuhnert et al. 2010; Kramer et al. 2011; Geier et al. 2015b; Geier and Lehnertz 2017; Baud et al. 2018; Campo et al. 2018). To our knowledge, our study is the first to investigate the long-term properties of functional brain networks and their correlation with seizure onset using scalp EEG.

Our results are in general agreement with (Kuhnert et al. 2010), who used iEEG data from 13 patients to monitor the long-term properties (characteristic path length, clustering coefficient) of binary functional brain networks constructed using mean phase coherence and thresholding to keep the mean degree constant across consecutive windows. A prominent 24-h rhythm, as well as shorter time periodicities were revealed in the temporal structure of the examined network measures. Also, similarly to our results, the effects of seizures on the latter were found to be considerably smaller in amplitude compared to the effect of physiological slower fluctuations. However, the shorter time periodicities and the correlation between longer term network fluctuations and seizure onset were not examined. In a subsequent study (Geier et al. 2015b), used a similar methodology to investigate the long-term evolution of degree-degree correlations (assortativity) in functional brain networks using iEEG data from seven patients suffering from pharmacoresistant focal epilepsy. Similarly, to our study, large fluctuations in time-resolved degree-degree correlations, which exhibited periodic temporal structure largely attributed to daily rhythms, were reported. Also, possible pre-seizure alterations were found to contribute marginally to the observed long-term fluctuations. The temporal and spatial variability of the importance of different regions in epileptic brain networks were investigated in (Geier and Lehnertz 2017) using iEEG data from 17 patients to construct networks using mean phase coherence. The importance of network nodes was assessed using strength centrality and betweenness centrality, which were subsequently used to define the corresponding important regions as the ones in which important nodes belong to. The importance of brain regions was found to fluctuate over time, with the fluctuations mostly attributed to processes acting on timescales of hours to days, with a strong contribution of daily rhythms.

In addition to the aforementioned studies, which examined the time evolution of summative network measures only, we demonstrated that the network topology is also characterized by the same periodic structure using a novel measure based on the graph edit distance (Eq. (9), Figures 4 and A.4). Our results revealed several additional periodicities at shorter time scales for all subjects both for the network summative properties (Fig. 3) and topology (Figs. 4, A.5). The most consistent of these shorter periodicities were harmonics of the main circadian periodicity, which were observed for all subjects (Figs. 1-2 and Table 2). In addition to these, subject-specific additional peaks in the PSD of the examined network properties were observed (e.g. Fig. 3). While these could be also due to physiological fluctuations, it cannot be ruled out that they are due to the reconfiguration of the underlying brain networks due to cognitive processes, which has been suggested to occur over much faster time scales – termed brain micro-states (Van De Ville et al. 2010).

Our results suggest that there is a significant correlation between the periodicities in network properties, particularly for the shorter consistently observed periodicities at 3.6 and 5.4 h, and seizure onset (Fig. 6, Tables 3,4). These results extend our previous work (Anastasiadou et al. 2016) and are in agreement with a recent study (Baud et al. 2018), which also used circular statistics to investigate correlations between seizure onset and brain dynamics using multi-day iEEG data from 37 subjects. Specifically, they obtained an epileptiform discharge measure on an hourly basis and used the wavelet transform to conclude that seizures were tended to occur during the rising phase of subject-specific, multi-dien rhythms. They also observed that the phase concentration was tighter for multi-dien rhythms as compared to circadian rhythms (Baud et al. 2018). In another recent study, (Karoly et al. 2017) investigated the time of seizure occurrence using long iEEG records (total of over 3,500 days) from a cohort of nine subjects. They too concluded that seizures tended to occur during preferred times of the day on a subject-specific basis and that incorporating this information can improve seizure prediction, using a logistic regression classifier.

Our results demonstrate the aforementioned correlations between long-term rhythms and seizure occurrence can be captured using scalp EEG, suggesting that it is not necessary to use an invasive modality (iEEG). The fact that the circadian rhythm is correlated to seizure onset suggests that seizures tend to agree at specific times for different subjects (Spencer et al. 2016; Karoly et al. 2017; Baud et al. 2018). However, the additional fact that seizure onset was more tightly correlated to the harmonics of the circadian periodicity suggests that in some cases, seizures do not occur not at the preferred time zone corresponding to the circadian periodicity, but at times separated by multiples of the period of these harmonic components (i.e. multiples of around 3.6 or 5.4 h). Importantly, the EEG signal power was not correlated to seizures (Figure 6), suggesting that network-based measures are a more sensitive marker of seizure occurrence. As seizure-induced changes in the network properties are considerably smaller in amplitude compared to longer-term rhythms (Figure 5), taking into account the instantaneous phase of the network-based periodicities can improve the sensitivity and specificity of seizure detection and prediction algorithms. For instance, detection or prediction based solely on constant, prespecified thresholds are not sufficient and are expected to yield either many false positives or negatives -if the selected threshold is relatively low or high respectively. It is also important to note that the examined network measures do not depend on selecting specific electrode locations.

### Study limitations

To account for the relatively low number of subjects, which is due to the practical difficulties of collecting long-term scalp EEG data, we applied multiple comparison corrections to our circular statistics results. However, note that previous related studies were based on similar subjects, e.g. (Kuhnert et al. 2010; Baud et al. 2018). To make sure that our analysis were not biased by the well-known EEG volume conduction effects (Nunez and Srinivasan 2006), we used the bipolar montage (Christodoulakis et al. 2013) and considered three correlation measures (correlation, corrected cross-correlation and coherence) that are differentially sensitive to volume conduction and reference choice effects. In general, while zero-lag correlations could be due to both artefactual (volume conduction/reference effects) and true correlations, non-zero lag correlations are more likely to reflect true correlations of underlying sources (Stam et al. 2007). By quantifying correlations by measures that are less sensitive to volume conduction, such as corrected cross-correlation, one accepts the risk of missing functionally meaningful correlations at zero-lag, but at the same time, the most frequent artefacts for misinterpretation of correlations are very much reduced (Stam et al. 2007). In the present case, the obtained results were similar for all measures, suggesting that volume conduction was not a major factor. Finally, the relation of seizures to sleep stages has also been examined, with previous studies suggesting e.g. that seizure rate was higher in non-REM sleep compared to REM sleep and demonstrated that on the descent from lighter into deeper levels of sleep, seizures are more likely to occur (Minecan et al. (2002)). We did not consider the effect of sleep staging here and we leave it to future studies.

## Acknowledgements

This work was partially supported by the European Regional Development Fund and the Republic of Cyprus through the Research Promotion Foundation (Project YΓEIA/ΔYΓEIA/0609(BE)/11).

